# Bona fide hematopoietic stem cells in zebrafish originate from the supra-intestinal artery

**DOI:** 10.1101/2025.05.20.655066

**Authors:** Shachuan Feng, Rui Qu, Yingzhu He, Tingting Huang, Youqi Wang, Thi Huong Trinh, Kefan Cheng, Shizheng Zhao, Ying Huang, Hao Zhu, Sicong He, Zilong Wen

**Author notes:** Correspondence should be addressed to ZLW: Department of Immunology and Microbiology, School of Life Science, Southern University of Science and Technology, Shenzhen 518055, China. S.F. and R.Q. contribute equally to this work.

## Abstract

Hematopoietic stem cells (HSCs) in vertebrates are widely believed to originate from hemogenic endothelial cells (HECs) in the dorsal aorta (DA) through endothelial-to-hematopoietic transition. However, conclusive in vivo DA-specific lineage-tracing studies to confirm this origin are lacking. Here, leveraging infrared light-mediated temporospatial-resolution lineage-tracing technology, we demonstrate that in zebrafish, HSCs do not originate from the DA. Instead, following the DA hematopoiesis, which produces hematopoietic precursors with limited renewal potential, lifelong renewable HSCs emerge from the supra-intestinal artery (SIA). Remarkably, while the DA produces hematopoietic cells from unipotent HECs, the SIA generates hematopoietic cells from both unipotent HECs and bipotent vessel-resident hemangioblasts (VRHs). These VRHs undergo multiple rounds of division and budding, generating both hematopoietic cells and SIA-resident endothelial cells. The distinct characteristics of DA and SIA cells are further reinforced by single-cell transcriptomic profiling. Our findings challenge the current paradigm of HSC development, underscoring the complexity of vertebrate hematopoiesis.

## INTRODUCTION

In vertebrates, most mature blood cells have short lifespans and must be continuously replenished by hematopoietic stem cells (HSCs), which possess the unique ability to self-renew and differentiate into all blood lineages throughout an organism’s lifespan.^1^ The development, maintenance, and differentiation of HSCs are governed by tightly regulated processes, and disruptions to these mechanisms are frequently linked to the pathogenesis of diverse diseases, including cancers.^2,3^ The importance of HSCs is further exemplified by their pivotal role in modern medical interventions, as HSC transplantation has emerged as a highly effective therapy for a wide array of severe blood disorders.^4,5^ Hence, elucidating the cellular and molecular mechanisms underlying HSC establishment and maintenance holds promise for advancing novel therapeutic strategies to treat blood-related diseases.

It has been well-accepted that HSCs in vertebrates are generated from the hemogenic endothelial cells (HECs) in the dorsal aorta (DA) through a process called endothelial-to-hematopoietic transitions (EHT).^6–9^ This process has been extensively studied and characterized in multiple vertebrate species, especially in mice and zebrafish. The first critical evidence supporting this concept came from the identification of transplantable HSC activity in the aorta-gonad-mesonephros (AGM) region of mouse embryos at embryonic day 9.5-10 (E9.5-10), as well as in ex vivo cultured AGM explants.^10–12^ These findings suggest that the AGM serves as the primary site of HSC generation in mice. Similarly, studies in zebrafish have revealed robust hematopoietic activity in the AGM, as evidenced by the strong expression of hematopoietic stem/progenitor cell (HSPC) markers in this region.^13–16^ Subsequent in vivo lineage tracing studies conducted in vasculature-specific Cre/CreER^T2^ reporter lines during embryonic stages have demonstrated that labeled cells give rise to various blood lineages that persist into adulthood in both mice and zebrafish,^7,17^ pointing to the endothelial origin of HSCs. As the DA is a prominent vascular structure associated with hematopoietic activity in the AGM of both mice and zebrafish,^6,12,18^ these studies collectively point to the DA as the principal source of HSC generation. This conclusion is further reinforced by the detection of hematopoietic cell clusters attached to the DA lumen in E9.5-10.5 mouse embryos and the visualization of EHT in the DA using ex vivo cultured AGM explants and live zebrafish embryos.^6–8,19,20^ Perhaps the most compelling evidence arises from the temporospatially controlled lineage-tracing experiments in zebrafish using the infrared light-evoked gene operator (IR-LEGO) system.^21–23^ These studies demonstrated that infrared light illumination of the AGM region housing the DA, prior to the onset of EHT, effectively labels multiple blood lineages that persist into adulthood.^21^ Collectively, these findings, along with additional supporting studies,^24–27^ have established the critical role of the DA in the HSC generation.

Despite these insightful studies, in vivo lineage tracing specifically targeting the DA endothelium to definitively confirm the DA as the primary origin of HSCs has been lacking. Recent research has indicated that lymphoid-biased fetal HSC-like cells, despite not naturally contributing to the adult HSC pool, can be reprogrammed into adult HSCs upon transplantation into irradiated recipient mice.^28^ This raises concern about the conclusions drawn from previous transplantation studies regarding the origin of HSCs. Additionally, several studies in mice have indicated the detection of pre-HSCs in the subaortic mesenchyme before the initiation of EHT in the DA, suggesting the possible existence of alternative sources for HSC generation.^29,30^ Furthermore, recent findings have revealed that the DA appears to be capable of producing non-HSC hematopoietic progenitors, including embryonic multipotent progenitors in mice and endothelium-derived hematopoietic progenitors (EHPs) in zebrafish,^21,31–34^ These discoveries highlight the complexity of hematopoiesis within this vascular structure. Finally, although temporospatially controlled lineage-tracing in zebrafish with the IR-LEGO system has shown that HSCs indeed originate from the AGM region,^21,23^ the resolution of the IR-LEGO system employed in these investigations is insufficient to distinguish the DA from other nearby vascular structures within the AGM. Therefore, there is a need for advanced lineage tracing methodologies with enhanced spatial resolution to accurately map the origin of HSCs.

The study described here employed a high-resolution IR-LEGO lineage-tracing system, combined with photoconvertible cell labeling and time-lapse live imaging, to investigate the origin of HSCs in zebrafish. The results reveal that HSCs in zebrafish originate from the supra-intestinal artery (SIA) rather than the DA. In contrast, the DA primarily generates EHPs that do not persist into adulthood. Through in vivo single-cell lineage tracing analysis, we discovered that, unlike the DA, where hematopoietic cells are generated from unipotent HECs, the SIA produces hematopoietic cells through both unipotent HECs and bipotent precursors identified as vessel-resident hemangioblasts (VRHs). These VRHs undergo multiple rounds of division and budding, giving rise to both hematopoietic and endothelial lineages. Finally, single-cell RNA sequencing (scRNA-seq) analysis unveiled distinct transcriptomic profiles and identified unique sets of transcription factors in cells originating from the DA and SIA. These findings reshape the foundations of HSC development studies, highlighting the intricate and multifaceted nature of vertebrate hematopoiesis.

## RESULTS

### Temporospatially restricted fate mapping challenges the dorsal aorta origin of HSCs

It has been documented that HSCs in vertebrates, including zebrafish, are generated from the HECs in the DA.^35^ Surprisingly, recent research has unveiled that zebrafish DA HECs also produce non-HSC hematopoietic progenitors, termed EHPs that are capable of giving rise to various blood lineages during larval and juvenile stages but not in adulthood.^21^ This raises questions about the mechanisms underlying the fate specification of HSCs versus EHPs in the DA. To investigate which HECs generate HSCs, we used a single-cell resolution IR-LEGO lineage-tracing system to label individual HECs in the DA and tracked their contribution to adult hematopoietic lineages (Figure 1A).^33^ To enhance the efficiency and precision of labeling, we developed a fate mapping line, *Tg(hsp70l:mCherry-T2A-CreER^T2^;kdrl:loxP-DsRedx-loxP-eGFP;coro1a:loxP-DsRedx-loxP-eGFP)* (*loxP-DsRedx-loxP-eGFP* is abbreviated to *LRLG* hereafter), featuring three cassettes: *hsp70l*-CreER^T2^ for infrared light inducible CreER^T2^ expression,^22,36^ endothelium-specific *kdrl-LRLG* for targeting DA HECs (*kdrl:*DsRedx^+^) and identifying embryos with successful labeling of a single HEC (*kdrl:*eGFP^+^), and hematopoiesis-specific *coro1a*-*LRLG* for visualizing hematopoietic derivatives (*coro1a:*eGFP^+^) (Figure 1A).^23,37^ Given that HSCs have been shown to originate predominantly from the anterior region of the DA,^21^ we focused on the anterior region (between the 8^th^ and 10^th^ somite) of the DA and illuminated a single HEC at 30 hours post-fertilization (hpf), prior to the onset of EHT.^6^ 10 hours post-IR illumination, embryos harboring a single *kdrl:*eGFP^+^ HEC in the ventral side of the DA were selected (Figures 1A and 1B) and raised to adulthood for flow cytometry analysis of adult hematopoietic organ—the kidney marrow (KM) (zebrafish counterpart of mammalian bone marrow)^38^ (Figures S1A and S1B). Surprisingly, only 4.4% (6 out of 136) of the treated fish exhibited successful labeling of HSCs, as evidenced by the presence of *coro1a:*eGFP^+^ hematopoietic cells encompassing lymphoid, myeloid, and hematopoietic precursor lineages in adult KM (Figures 1C and S1B). This percentage falls significantly below the estimated HSC frequency (∼60%, corresponding to ∼21 HSC precursors and ∼34 DA floor HECs)^39,40^ and is even lower than the control fish (7.5%, 20/268) (Figure 1C), indicating that the detection of hematopoietic lineages in adult KM in both control and DA-targeted groups is caused by sporadic leaky activity of CreER^T2^. Given that the single cell labeling specifically targeted the anterior DA (between 8^th^ and 10^th^ somite) where HSCs are presumed to be most-enriched,^21^ these results suggest that the DA contributes minimally, if at all, to HSC generation.

**Figure 1.**
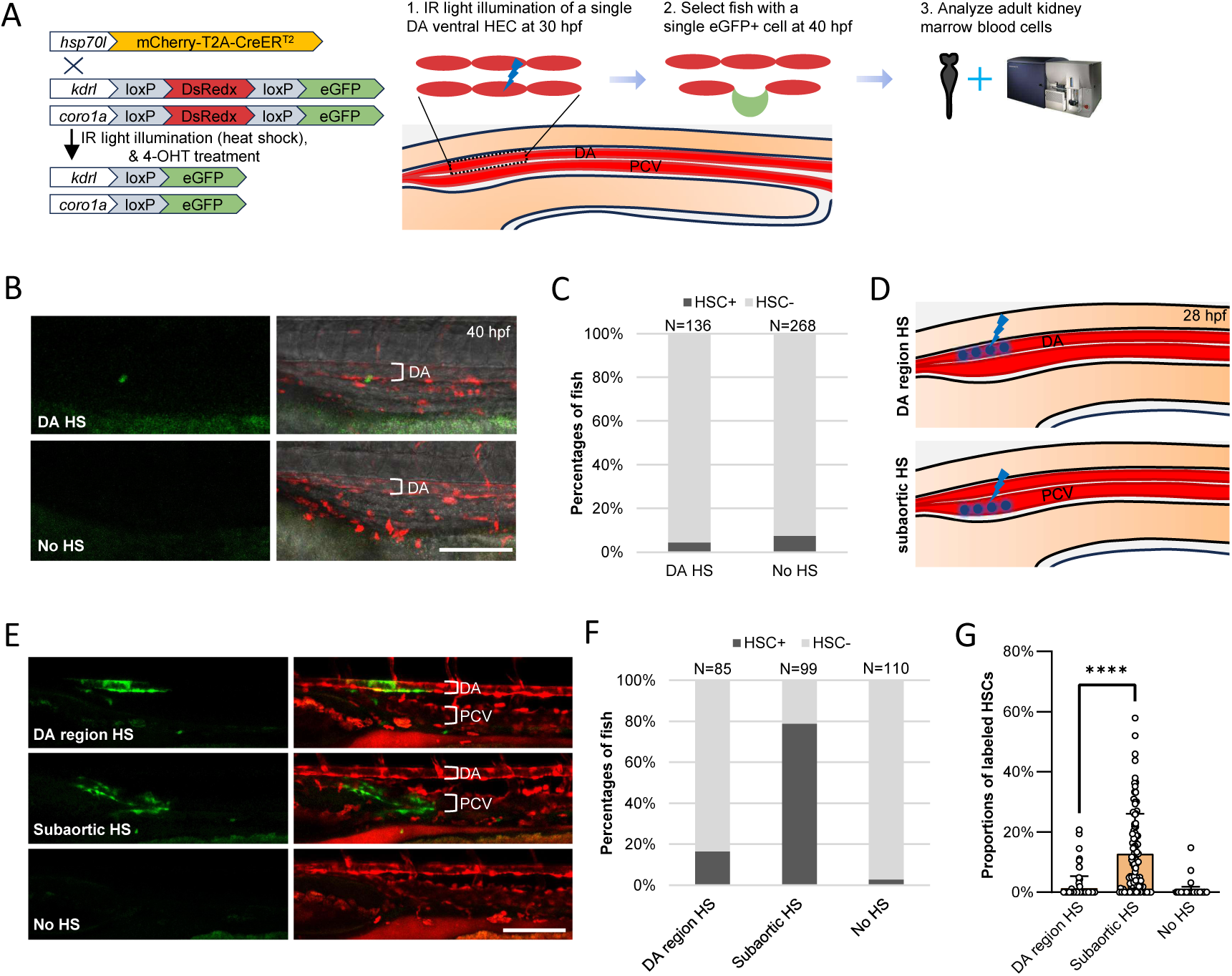
HSCs in zebrafish do not originate from the DA. (A) The left panel illustrates the principle of the single-cell resolution IR-LEGO lineage tracing strategy. In 30 hpf *Tg(hsp70l:mCherry-T2A-CreER^T2^;kdrl:LRLG;coro1a:LRLG)* embryos, a single HEC is illuminated by infrared light (heat shock) followed by 4-OHT treatment, inducing CreER^T2^-mediated recombination of the reporter genes. This results in the targeted single HEC and its hematopoietic progeny switching from DsRedx to eGFP expression. The right panel outlines the experimental pipeline. DA: dorsal aorta. PCV: posterior cardinal vein. (B) Representative images of a successful single cell-labeled fish (upper panel) and a control fish (lower panel). 10 hours post-IR illumination, fish with single HEC labeling (eGFP^+^) in the DA, confirmed by confocal microscopy, were selected for further analysis. Scale bar: 100 µm (all scale bars hereafter are 100 µm unless otherwise annotated). HS: heat shock. (C) Summary of the ratios of HSC-labeled fish in the single cell-labeled (DA HS) and control (No HS) groups, based on flow cytometry analysis of adult kidney marrow blood cells (3 to 12 months fish) (Fig. S1A and S1B). 136 DA HS fish (N=136) and 268 control fish (N=268) were evaluated. The data is a combination of 22 independent experiments. In each experiment, 4-8 single cell-labeled fish were obtained from 20-30 infrared light-illuminated fish. (D) Schematic illustration of multi-cell resolution infrared light-mediated labeling of one somite-width DA region or subaortic AGM region between 8^th^ and 10^th^ somite. (E) Representative images of DA-labeled, subaortic AGM region-labeled, and control (No HS) fish. Fish with relatively region-specific eGFP^+^ vasculature pattern observed via confocal microscopy on 3 dpf were selected and survived to adulthood for further analysis. (F) The ratios of HSC-labeled fish in the three groups. The results are derived from a combination of 11 independent experiments to ensure sufficient sample size for reliable percentages. (G) The proportions of labeled adult HSCs (defined by the proportions of labeled KM myeloid lineages) in the three groups show significant differences. The subaortic AGM region-targeted group displays a significantly higher mean efficiency (12.8%) compared to the DA region-targeted group (1.35%) and the control (No HS) group (0.23%), with the highest labeling efficiency of around 60%. Data are presented as Mean ± SD. Welch’s t test was used for significance calculations. **** P<0.0001.

### HSC precursors are spatially restricted to the subaortic AGM region

While previous multi-cell resolution IR-LEGO studies have confirmed HSC emergence from the AGM region,^21,23^ our single-cell labeling data suggest that these HSCs likely originate from non-DA tissues within this domain. We thus proposed the subaortic AGM region, where two key components, the gonad and mesonephros of the AGM, are located,^41–43^ as the most probable HSC source.

To support this speculation, we applied multi-cell resolution IR-LEGO system to label one somite width region between 8^th^ and 10^th^ somite targeting the anterior subaortic AGM region at 28 hpf (Figure 1D) and then tracked their contributions to hematopoietic lineages in adult KM. In parallel, we also targeted the adjacent anterior segment of the DA as the control. The labeling specificity was confirmed by confocal imaging of *kdrl*:eGFP signals in both the subaortic posterior cardinal vein (PCV) and the DA, and embryos with minimal cross-labeling were raised to adulthood for flow cytometry analysis of KM blood cells (Figure 1E). As anticipated, 79% (78 out of 99) of the subaortic AGM region-targeted fish displayed abundant *coro1a:*eGFP^+^ hematopoietic cells, encompassing lymphoid, myeloid, and hematopoietic precursor lineages, in adult KM (Figures 1F and S1C). In contrast, only 16% (14 out of 85) of the DA region-targeted fish showed coro1a:eGFP+ hematopoietic cells in adult KM (Figures 1F and S1C), a moderate increase compared to the untargeted fish (2.7%, 3 out of 110). More importantly, an average of 12.8% of adult KM HSCs were labeled in the subaortic AGM region-targeted fish, a striking contrast to 1.35% in the DA region-targeted fish and 0.23% in control fish (Figure 1G). The minimal labeling percentage of hematopoietic cells in the DA region-targeted fish suggests that the observed HSC labeling in this group is most likely due to sporadic CreER^T2^ leaky activity and unavoidable occasional cross-labeling of the subaortic AGM region by multi-cell resolution IR illumination. Collectively, these results, combined with single-cell fate mapping data, suggest that the majority of HSCs do not originate from the DA but instead arise from precursor cells residing in the subaortic AGM region.

### HSCs arise from previously unidentified precursors located beneath the anterior region of the PCV

Previous studies in both mammals and zebrafish have supported the endothelial origin of HSCs.^21,23^ As PCV is the major vasculature in the subaortic AGM region,^44^ we hypothesized that PCV might be the source giving rise to HSCs. We therefore performed time-lapse live imaging on venous endothelium reporter line *Tg(stab1:DsRedx)* to determine whether EHT occurs in the PCV. Surprisingly, no EHT was observed during the imaging period spanning from 29 hpf to 72 hpf (Figure 2A; Movie S1), indicating that HSCs are unlikely to originate directly from the PCV. To reinforce this notion, we crossed the lymphatic/venous endothelium-specific CreER^T2^ line *Tg(lyve1b:CreER^T2^)*^45^ with the *Tg(kdrl:LRLG;coro1a:LRLG)* reporter line to generate triple transgenic fish for fate mapping analysis (Figure 2B). As predicted, despite effectively labeling nearly all PCV endothelial cells (Figure 2B), only 8% (6 out of 71) of the fish showed limited *coro1a*:eGFP^+^ hematopoietic cells in adult KM, with an average HSC-labeling efficiency of only 0.61% (Figures 2C and 2D). In contrast, all pan-endothelial *Tg(kdrl:CreER^T2^)* fish (15 out of 15) exhibited robust HSC-labeling, with an average HSC-labeling efficiency of 66.97% (Figures 2C, 2D, and S2A). These findings, combined with time-lapse live imaging data, confirm that HSCs do not originate from the PCV but most likely arise from undisclosed *kdrl*-positive precursors situated in the vicinity of the anterior PCV.

**Figure 2.**
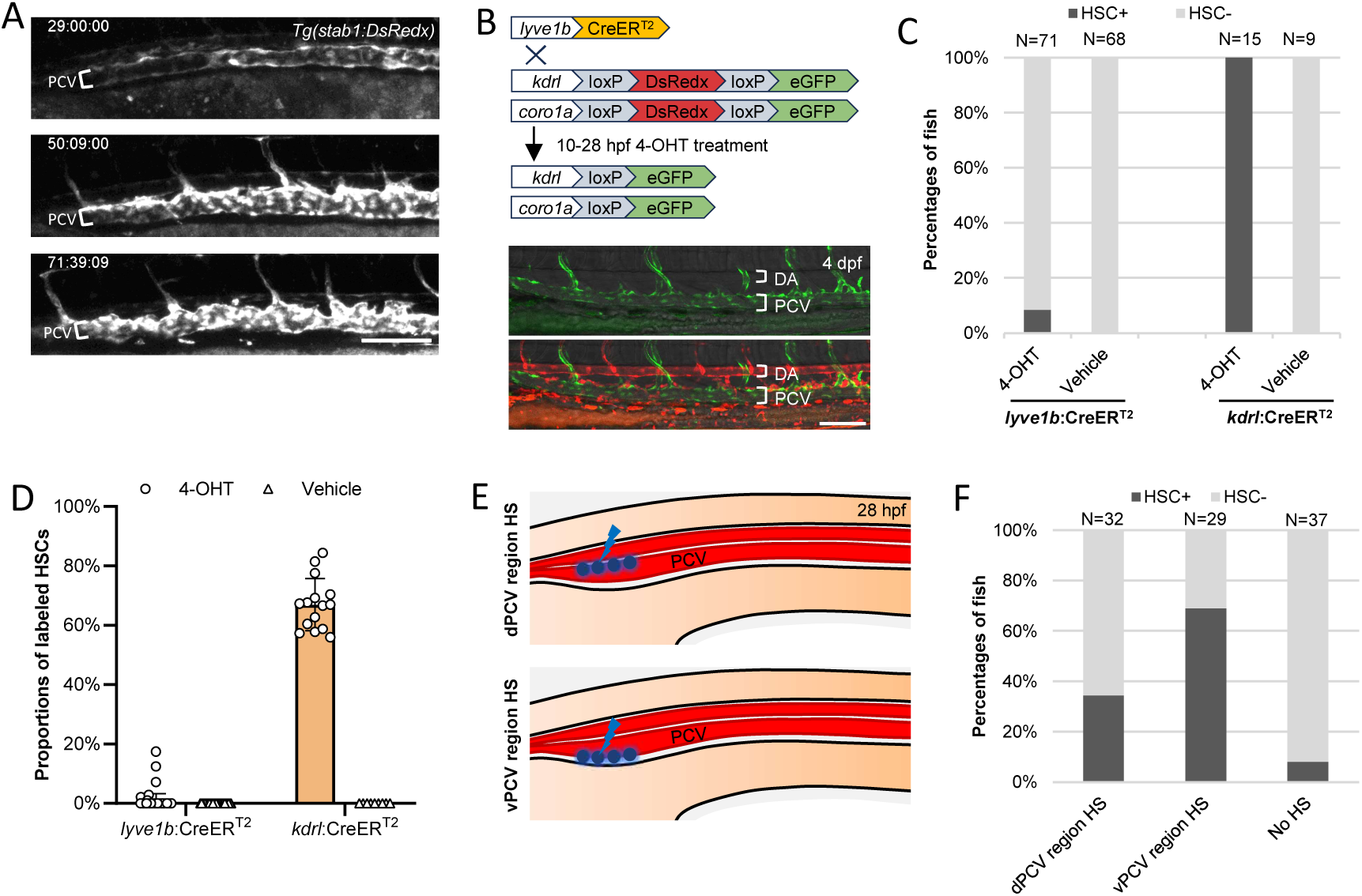
The posterior cardinal vein does not generate HSCs. (A) Representative images of time-lapse live imaging of anterior PCV during 29 hpf to 72 hpf in the *Tg(stab1:DsRedx)* fish. EHT-like events have not been detected in over 10 examined fish; only some autofluorescence-positive pigment cells are visible in the subvenous region. (B-D) Venous vessel-specific lineage tracing denies the PCV as the origin of HSCs. The experimental principle is elucidated in the schematic diagram (B, upper panel). Fish were treated with 4-OHT from 10 hpf to 28 hpf, and labeling efficiency and specificity were confirmed by the eGFP^+^ vasculature pattern at 3 dpf using confocal microscopy (B, lower panel). Flow cytometry analysis of adult KM blood cells revealed that fewer than 9% of the *lyve1b*:CreER^T2^-activated fish (6/71) showed eGFP^+^ blood cells in adult KM (C, left two columns), with an average of 0.61% HSC-labeling efficiency (D, left column). In contrast, all pan-vessel-specific *kdrl*:CreER^T2^-activated fish (positive control) display eGFP^+^ blood cells in adult KM (C, right two columns), with an average of 66.97% HSC-labeling efficiency (D, right column). To ensure sufficient sample size for reliable percentages, the *lyve1b*:CreER^T2^-mediated and *kdrl*:CreER^T2^-mediated lineage tracing data combine results from 4 and 2 independent experiments, respectively. (E-F) Multi-cell resolution IR-LEGO lineage tracing indicates that HSCs arise from unidentified precursors located beneath the PCV. The schematic draw indicates the positions (dorsal and ventral sides of the PCV) targeted by infrared light in *Tg(hsp70l:mCherry-T2A-CreER^T2^;kdrl:LRLG;coro1a:LRLG)* fish at ∼28 hpf (E). Flow cytometry analysis of adult KM blood cells reveals a significant higher percentage of HSC-labeled fish in the ventral PCV region-targeted group (vPCV) compared to the dorsal PCV region-targeted group (dPCV) (F). The data represents a combination of three independent experiments, ensuring sufficient sample size for reliable percentages.

To further identify the precise location of these undisclosed precursors, we applied the IR-LEGO lineage-tracing system to target either the dorsal or ventral side of the anterior PCV in the *Tg(hsp70l:mCherry-T2A-CreER^T2^;kdrl:LRLG;coro1a:LRLG)* reporter line and assessed HSC-labeling frequency in each group (Figures 2E and S2B). The results showed that ∼70% of the fish targeted on the ventral side contained abundant *coro1a:*eGFP^+^ hematopoietic cells in adult KM, compared to only 34% in the dorsal-targeted group (Figure 2F). Together, these findings indicate that the majority of undisclosed HSC precursors are located beneath the anterior PCV.

### The supra-intestinal artery gives rise to HSCs via EHT

Given the fact that most adult zebrafish hematopoietic cells are likely of endothelial origin,^7^ we hypothesized that HSCs are probably derived from a subset of endothelial precursors positioned beneath the anterior PCV. Notably, three vascular structures are known to develop beneath the PCV during zebrafish embryogenesis: the supra-intestinal artery (SIA), sub-intestinal vein (SIV), and sub-intestinal venous plexus (SIVP).^44,46–49^ We reasoned that one or more of these vascular structures might serve as the source of HSCs. To test this, we examined these vessels from 2 days post fertilization (dpf) to 7 dpf, after the SIA, SIV, and SIVP are well-established, for potential budding cells. Confocal imaging in *Tg(fli1:eGFP)* fish did not capture any potential budding cells before 4 dpf (Figure S3A), but by 7 dpf, a few potential budding cells were detected in the anterior SIA but not in the SIV and SIVP (Figure S3B). This suggests that the SIA may be the source generating HSCs from 4 dpf onwards. To support this speculation, we conducted time-lapse live imaging using the *Tg(kdrl:eGFP)* reporter line from 4 dpf onwards. Remarkably, we found that the endothelium in the anterior SIA, but not in the posterior SIA, the SIV, and the SIVP, undergoes EHT, producing cells resembling hematopoietic cells (Figure 3A; Movie S2). To verify that SIA-derived cells are indeed hematopoietic cells, we photoconverted the anterior SIA in 4 dpf *Tg(kdrl:Dendra2)* larvae and tracked the fates of the converted cells at later stages. The results showed that the photoconverted Dendra2-red cells were increasingly detected in the thymus and caudal hematopoietic tissue (CHT), two well-known hematopoietic tissues in zebrafish,^50,51^ from 5 dpf to 7 dpf, documenting their hematopoietic identity (Figures 3B and 3C).

**Figure 3.**
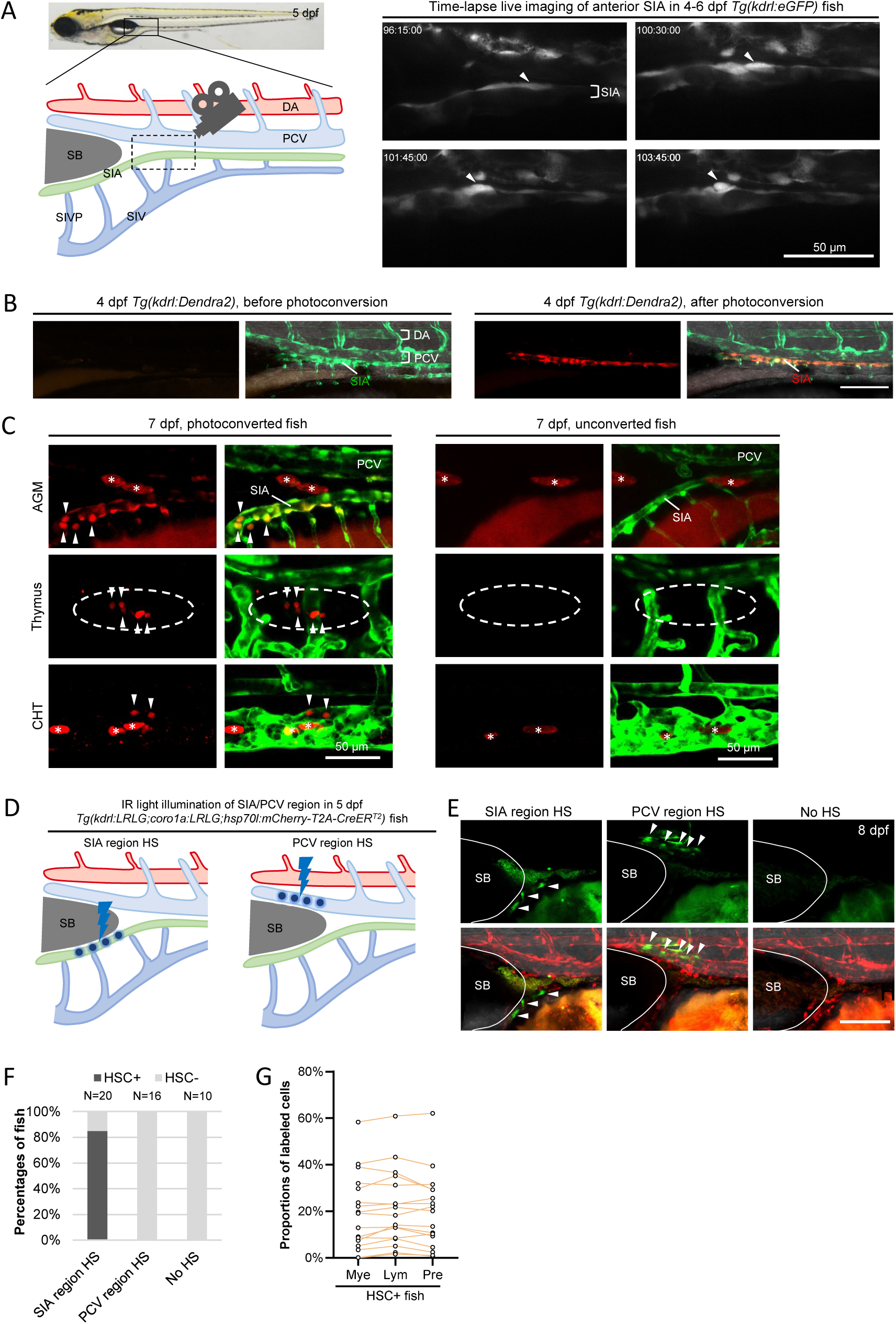
HSCs are generated from the supraintestinal artery via EHT. (A) Time-lapse live imaging of the subvenous vasculatures in 4 dpf to 6 dpf *Tg(kdrl:eGFP)* fish reveals EHT-like events in the SIA. The left panel depicts subvenous vasculature structures in the anterior AGM region, while the right panel shows a representative EHT-like event (arrowhead) in the SIA, originating from the dashed line-boxed region in the left panel. SIA: supra-intestinal artery. SIV: sub-intestinal vein. SIVP: sub-intestinal venous plexus. SB: swimming bladder. (B-C) Photoconversion-based transient lineage tracing reveals SIA-derived cells colonizing the thymus and CHT. Representative images of 4 dpf *Tg(kdrl:Dendra2)* fish indicate anterior SIA cells before and after the photoconversion (B). The photoconverted SIA cells give rise to hematopoietic cells-like descendants (arrowheads) that colonize the AGM, thymus (dished line circle), and CHT (C). Autofluorescence-positive pigment cells are marked by asterisks. (D-G) The long-term lineage tracing using multi-cell resolution IR-LEGO cell labeling confirms the SIA as the origin of HSCs. The schematic diagram (D) indicates the anterior PCV region and SIA region targeted by infrared light in *Tg(hsp70l:mCherry-T2A-CreER^T2^;kdrl:LRLG;coro1a:LRLG)* zebrafish at 5 dpf. Labeling specificity was validated by eGFP^+^ vasculature pattern three days post-infrared light illumination (E). Flow cytometry analysis of the adult KM blood cells reveals successful HSC labeling in 85% SIA region-targeted fish (17 out of 20), whereas no labeling was observed in the PCV region-targeted group and No HS (control) fish (F). Adult blood cell-labeling efficiency, determined by assessing the proportions of labeled KM blood lineages (eGFP^+^ cells) in all KM blood cells (DsRedx^+^ + eGFP^+^ cells) in each HSC-labeled fish, is shown in panel G. The data represents a combination of three independent experiments to ensure sufficient sample size for reliable percentages.

To confirm the presence of HSCs among SIA-derived hematopoietic cells, we employed muti-cell resolution IR-LEGO system to permanently label the anterior SIA region in the *Tg(hsp70l:mCherry-T2A-CreER^T2^;kdrl:LRLG;coro1a:LRLG)* reporter line at 5 dpf, a stage where the anterior SIA is well-separated from the PCV by the swimming bladder (Figure 3D).^52^ The anterior PCV region was also labeled as a control. The infrared light labeling of the SIA and PCV was highly specific, with no cross-labeling observed (Figure 3E). The IR-illuminated fish were raised to adulthood and analyzed by flow cytometry for adult KM (Figure S3C). Consistent with our hypothesis, 85% of SIA region-targeted fish exhibited abundant *coro1a:*eGFP^+^ hematopoietic cells (lymphoid, myeloid, and precursor lineages), while these cells were absent in the PCV region-targeted group (Figure 3F). Among the SIA region-targeted fish with successfully labeled adult blood cells, ∼20% of all three major cell populations were labeled on average, with the highest labeling efficiency reaching ∼60% (Figure 3G). Collectively, these findings, combined with time-lapse live imaging and photoconversion assays, demonstrate that the majority of HSCs are generated from the anterior SIA through EHT.

### HSC emergence from the supra-intestinal artery requires Runx1 and Notch signaling

The transcription factor RUNX1/Runx1 is known to play a crucial role in EHT in the DA of both mammals and zebrafish.^6,53^ We thus hypothesized that Runx1 also regulates EHT in the SIA. To test this, we crossed *runx1*-deficient *runx1^w84x^*mutants with the *Tg(kdrl:eGFP)* reporter line and quantified SIA-associated *kdrl:*eGFP^+^ cells,^54,55^ which likely represent nascent HSCs derived from the SIA (Figures 4A and 4B). As expected, *runx1^w84x^* mutants showed a significant reduction in SIA-associated *kdrl:*eGFP^+^ cells (Figure 4B), indicating that *runx1* deficiency disrupts the formation of SIA-born HSCs. To confirm this was due to EHT defects, we performed time-lapse live imaging on 4-6 dpf *Tg(kdrl:eGFP);runx1^w84x^* mutant larvae. Consistent with our hypothesis, EHT in the SIA was nearly absent in the mutants (Movie S3). Collectively, these findings demonstrate that, similar to the DA,^6^ *runx1* is essential for hematopoietic cell generation in the SIA.

**Figure 4.**
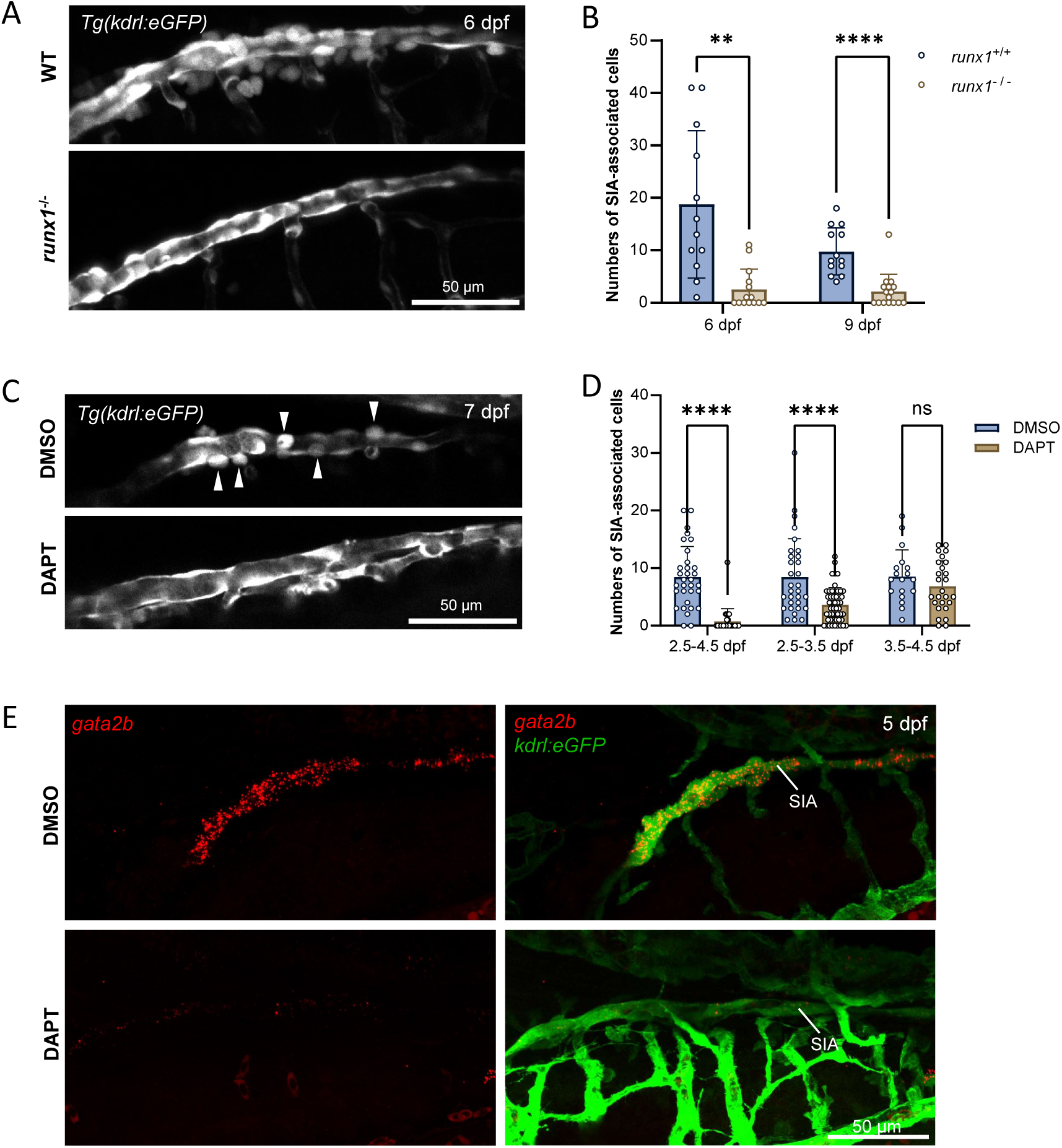
SIA hematopoiesis depends on *runx1* and Notch signaling. (A-B) SIA-associated HSPC-like cells are significantly reduced in *runx1*^-/-^ *Tg(kdrl:eGFP)* fish. Data are presented as Mean ± SD. Welch’s t test was utilized for significance calculations in 6 dpf samples, and Student’s t-test was used for 9 dpf samples. ** P<0.01. **** P<0.0001. (C-E) Notch inhibitor treatment blocks SIA hematopoiesis by suppressing HEC formation. Representative images and statistical data show SIA-associated HSPC-like cells (arrowheads) in 7 dpf DMSO-treated control fish, while these cells are largely absent in fish treated with DAPT from 2.5 dpf to 4.5 dpf (C, D). Additional DAPT treatment experiments confirm that Notch signaling is essential for SIA hematopoiesis before 3.5 dpf, prior to EHT onset (D), likely by disrupting HEC formation. The importance of Notch signaling is further supported by the lack of *gata2b* expression in the SIA, as shown by RNAscope in situ hybridization in 5 dpf *Tg(kdrl:eGFP)* fish treated with DAPT (from 2.5 dpf to 4.5 dpf) compared to the DMSO-treated controls (E). Data are presented as Mean ± SD. Welch’s t-test was used for significance calculations in the 2.5-4.5 dpf and 2.5-3.5 dpf treatment samples, while Student’s t-test was employed for the 3.5-4.5 dpf treatment samples. **** P<0.0001. ns: non-significant.

Previous studies have established the critical role of Notch signaling in HSPC formation in both mammals and zebrafish.^56–58^ However, the conclusions in zebrafish primarily rely on detecting *runx1^+^* HECs in the DA and *runx1^+^*/*cmyb^+^*/*cd41^+^*hematopoietic cells in the AGM region.^57,59^ which, as our fate-mapping study shows, are largely EHPs derived from the DA (Figures 1C, 1F, and 1G). Thus, it is essential to reassess the role of Notch signaling in the development of bona fide HSCs originating from the SIA. To address this, we treated the *Tg(kdrl:eGFP)* embryos with the Notch inhibitor DAPT^60^ from 2.5 dpf to 4.5 dpf, shortly after SIA formation,^46^ and evaluated its impact on SIA hematopoiesis by quantifying SIA-associated *kdrl:*eGFP^+^ HSPC-like cells a few days later. Indeed, DAPT treatment significantly reduced SIA-associated *kdrl:*eGFP^+^ cells (Figures 4C and 4D), underscoring the pivotal role of Notch signaling in SIA hematopoiesis. To pinpoint the stage at which Notch signaling acts, we administered DAPT treatment to embryos during two windows: 2.5-3.5 dpf (prior to EHT initiation) and 3.5-4.5 dpf (after EHT initiation). Only the treatment during 2.5-3.5 dpf significantly reduced SIA-associated *kdrl:*eGFP^+^ cells (Figures 4C and 4D). suggesting Notch signaling regulates SIA hematopoiesis before EHT initiation, likely by influencing the formation of the HECs. To validate this, we performed RNAscope in situ hybridization of *gata2b*, a well-known regulator of HEC fate,^61^ in DAPT- and DMSO-treated 5 dpf *Tg(kdrl:eGFP)* fish. As anticipated, *gata2b* was robustly expressed in the majority of anterior SIA cells in the DMSO-treated fish but largely absent in SIA in DAPT-treated fish (Figure 4E), consistent with the role of Notch signaling in HEC formation in the DA.^40,57,62^ Notably, in DMSO-treated fish, the majority of anterior SIA cells exhibited *gata2b* expression, further supporting the idea that the SIA lacks the dorsal-ventral polarity and that most, if not all, anterior SIA cells possess hemogenic potential.

### Supra-intestinal artery hematopoiesis involves vessel-resident hemangioblasts and sustains an extended period

To unveil the dynamics of HSC generation in the SIA, we leveraged the *Tg(kdrl:eGFP)* reporter line to quantify eGFP^+^ cells closely associated with the SIA, which likely represent nascent HSCs derived from the SIA, starting at 5 dpf (Figure 5A). We found that SIA-associated eGFP^+^ cells were consistently detected at 5 dpf, increased steadily to a peak of ∼15 cells by 7-8 dpf, and then gradually declined from 10 dpf onwards, remaining detectable until 15 dpf (Figure 5B).

**Figure 5.**
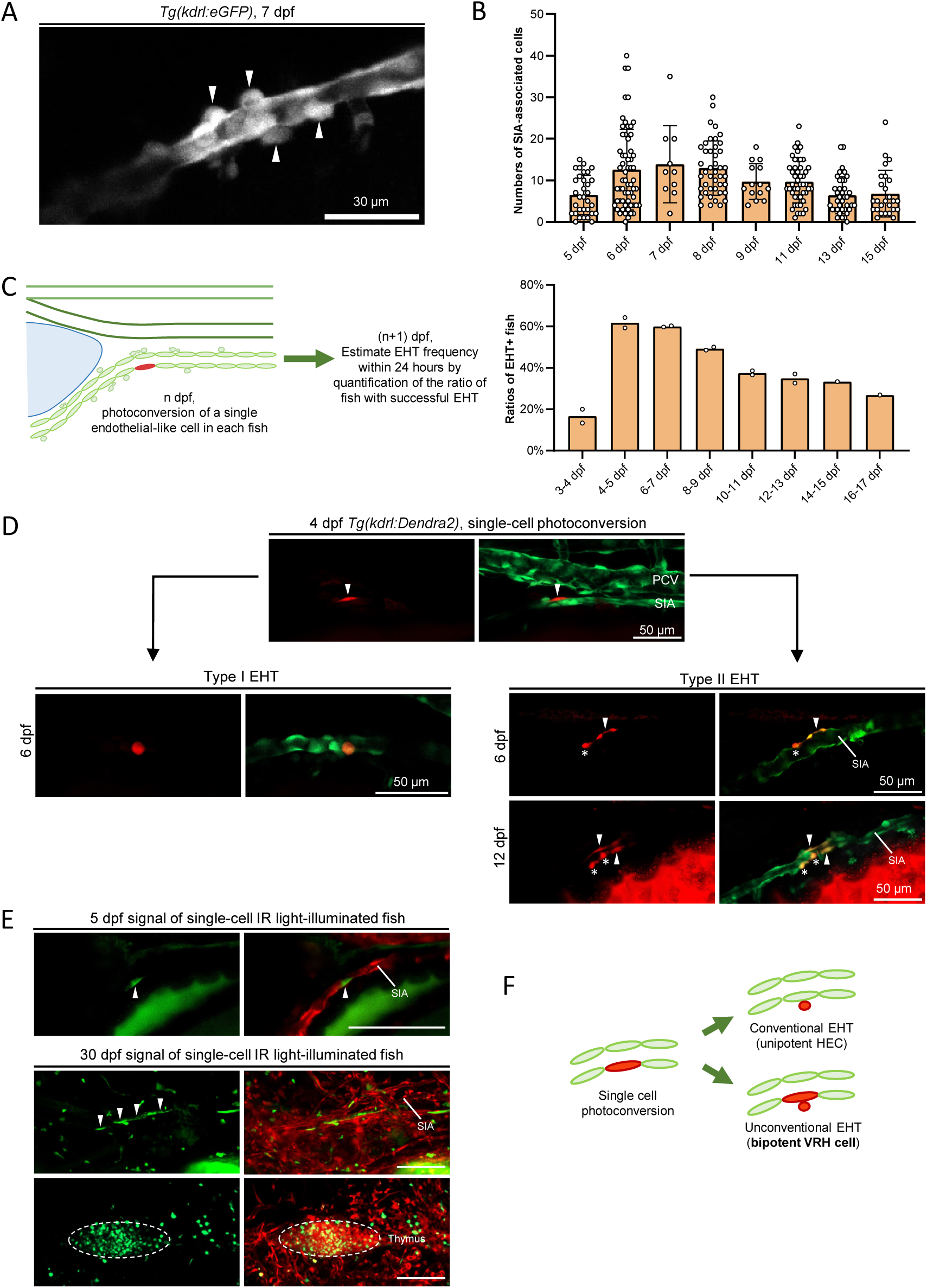
SIA hematopoiesis persists for at least two weeks through continuous cell division and budding of VRHs. (A-B) Characterization of SIA-associated nascent HSPC-like cells. A representative confocal microscopy image shows the SIA-associated nascent HSPC-like cells (arrowheads) (A). Quantification analysis reveals SIA-associated nascent HSPC-like cells persist from 5 dpf to at least 15 dpf (B). (C) Single-cell photoconversion assay in the *Tg(kdrl:Dendra2)* fish unveils SIA EHT dynamics. The schematic diagram illustrates experimental processes, with EHT events evaluated 24 hours-post photoconversion. Quantification analysis shows that SIA EHT initiates at ∼4 dpf, peaks at approximately 6 dpf, and gradually declines after 7 dpf. (D) Short-term lineage tracing of photoconverted individual SIA cells identifies two EHT processes. In one process, photoconverted cells directly bud out from the SIA, producing only HSPC-like cells. In the other, photoconverted cells undergo multiple rounds of division and budding, generating both hematopoietic daughter cells (asterisks) and endothelial daughter cells that remain in the SIA (arrowheads) until at least 12 dpf. This observation suggests that the SIA contains both unipotent conventional HECs and bipotent precursors termed vessel resident hemangioblasts (VRHs). (E) Long-term lineage tracing of infrared light-labeled individual SIA cells in *Tg(hsp70l:mCherry-T2A-CreER^T2^;fli1:LRLG;coro1a:LRLG)* zebrafish confirms the presence of VRHs. A successfully labeled single SIA cell (arrowhead) is shown in the upper panel. Subsequent tracing revealed abundant hematopoietic (including T lymphocytes in the thymus) and endothelial (arrowheads) descendants by 30 dpf (lower panels). (F) Schematic summary of the two EHT types in the SIA.

Since the number of SIA-associated eGFP^+^ cells reflects a combination of EHT along with the proliferation and migration rates of these cells, it may not accurately represent the true dynamics of SIA hematopoiesis. To better elucidate these dynamics, we utilized endothelium-specific *Tg(kdrl:Dendra2)* transgenic fish^21,63^ and photoconverted single HECs/ECs in the anterior SIA from 3 dpf to 16 dpf. 24 hours post-photoconversion, we assessed the EHT frequency by calculating the ratio of fish with SIA-associated Dendra2-Red cells, presumably representing the hematopoietic cells derived from converted HECs (Figure 5C). We found that SIA EHT commenced around 4 dpf, with ∼17% of converted cells undergoing EHT (Figure 5C). EHT frequency rapidly increased, peaking at ∼62% at 4-5 dpf, then gradually declined from 10-11 dpf onwards, stabilizing at a lower level (∼27%) until ∼17 dpf (Figure 5C). These findings indicate that, unlike the DA where the hematopoiesis initiates early in embryogenesis and diminishes within ∼2 days (30-68 hpf),^6^ SIA hematopoiesis emerges later during the larval stage and persists for at least two weeks.

Intriguingly, we observed that, in addition to generating hematopoietic progeny via canonical EHT, where photoconverted single cells directly bud out from the SIA (Figure 5D),^40^ a subset of converted cells underwent multiple rounds of division and budding, producing both hematopoietic daughter cells and endothelial daughter cells that remained within the SIA (Figure 5D). This prompted us to speculate that these SIA cells are bipotent precursors capable of giving rise to both hematopoietic and endothelial lineages. To support this, we employed the single-cell IR-LEGO lineage tracing system to permanently label a single cell in the anterior SIA of 4 dpf *Tg(hsp70l:mCherry-T2A-CreER^T2^;fli1:LRLG;coro1a:LRLG)* reporter fish as described previously^33^ and tracked the fate of the labeled SIA cells until 30 dpf, when SIA EHT near completion. Indeed, a significant number of labeled fish contained both *coro1a*:eGFP^+^ hematopoietic cells in various tissues and *fli1:*eGFP^+^ endothelial cells in the SIA (Figure 5E), strongly supporting the bipotential of these SIA cells, which we term as vessel-resident hemangioblasts (VRHs). Together, these findings demonstrate that the SIA comprise unipotent HECs and bipotent VRHs, which generate hematopoietic cells through canonical EHT and non-canonical EHT, respectively (Figure 5F).

### scRNA-seq analysis reveals transcriptomic signatures of cells in the supra-intestinal artery and dorsal aorta

To explore the transcriptomic characteristics of HECs and ECs in the SIA, we photoconverted the anterior SIA cells in 5 dpf *Tg(kdrl:Dendra2)* zebrafish and isolated the Dendra2-Red cells for scRNA-seq analysis. By combining this SIA scRNA-seq data with 28 hpf DA scRNA-seq data,^40^ we performed uniform manifold approximation and projection (UMAP) analysis. The result revealed four distinct clusters, each cluster containing cells from both DA and SIA cells, indicating a significant similarity between these two hematopoietic processes (Figures 6A and 6B). Based on endothelial and hematopoietic marker gene expression, we annotated the clusters as EC, HEC_1, HEC_2, and HSPC (Figures 6A and 6C). The EC cluster exhibited strong expression of endothelial feature genes (*etsrp*, *cdh5*, *kdrl*, *fli1*, *efnb2a*, *dll4*, *flt1*, and *hey2*), while the HEC and HSPC clusters showed pronounced hematopoietic potential, as indicated by the enrichment of hematopoietic markers (*gata2b*, *runx1*, *myb*, *spi2,* and *gfi1aa*,), which could be further distinguished by the differing expression of terminal hematopoietic genes including *coro1a*, *csf1rb*, and *ptprc* (Figure 6C). Notably, despite clear separation of the four clusters, the HEC_1 and HEC_2 populations displayed a high degree of similarity (Figure 6D), suggesting minimal transcriptomic differences between them. Therefore, we combined the two HEC populations for subsequent comparative analysis.

**Figure 6.**
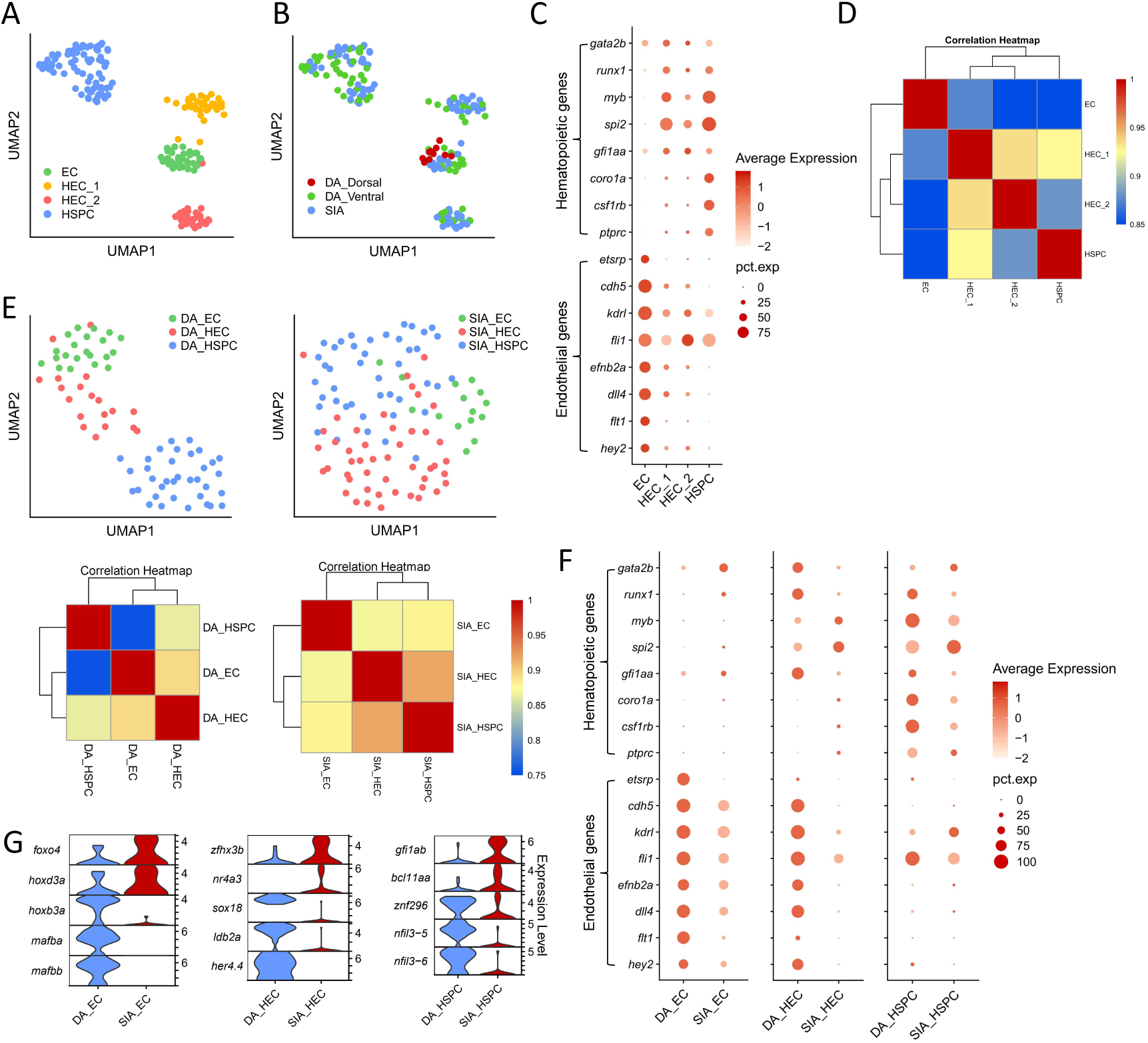
scRNA-seq supports the presence of VRHs and identifies potential regulators of SIA hematopoiesis. (A-B) UMAP analysis of 5 dpf SIA cells and 28 hpf DA cells reveals four distinct cell clusters (A) and their respective origins (B). (C) Dot plot illustrates the expression patterns of selected hematopoietic and endothelial marker genes in the four clusters, defining their identities. (D) Correlation heatmap reveals significant similarity between the HEC_1 and HEC_2 clusters. (E) UMAP analysis (upper panel) and correlation analysis (lower panel) of DA and SIA clusters (HEC_1 and HEC_2 are merged as HEC) highlight their distinct characteristics: DA clusters are well-separated with lower correlation, while SIA clusters are largely intermingled with high correlation. (F) Comparison of selected endothelial and hematopoietic marker expression in SIA and DA cells. DA ECs express strong endothelial markers but lack hematopoietic markers (left panel), while SIA ECs show weaker endothelial markers and express several notable hematopoietic markers like *gata2b*, *runx1*, and *gfi1aa* (left panel), supporting their bipotent nature. SIA HECs (middle panel) and SIA HSPCs (right panel) exhibit distinct hematopoietic marker gene expression compared to DA counterparts, suggesting unique regulatory mechanisms in SIA hematopoiesis. (G) Differential gene expression (DEG) analysis identifies distinct sets of transcription factors differentially expressed between SIA and DA cells.

To better understand the transcriptomic differences between the cells from the DA and those from the SIA, we subdivided the EC, HEC (merged HEC_1 and HEC_2 populations), and HSPC populations based on origins and performed UMAP analysis for each. Interestingly, while DA subpopulations were distinct and well-separated in the UMAP, SIA subpopulations were intermingled (Figure 6E), indicating that transcriptomic differences among SIA subpopulations are less pronounced compared to those between DA subpopulations. This observation aligns with the correlation analysis (Figure 6E) as well as our experimental data, which suggest that anterior SIA cells are hemogenic and lack the dorsal-ventral polarity seen in the DA (Figures 4E and 5A). Strikingly, analysis of endothelial and hematopoietic marker genes revealed that while DA_EC subpopulation strongly expressed endothelial markers but lacked hematopoietic markers, SIA_EC subpopulation not only expressed endothelial markers but also showed detectable levels of hematopoietic genes, particularly *gata2b*, a key regulator of HEC formation (Figure 6F).^61^ This suggests the potential bipotency of SIA_ECs, consistent with our lineage tracing data identifying VRHs (Figures 5D-5F). Additionally, SIA_HECs and SIA_HSPCs exhibited reduced expression of key hematopoietic genes (*gata2b*, *runx1*, and *gfi1aa*) compared to their DA counterparts (Figure 6F), highlighting intrinsic differences may underlie the distinct hematopoietic processes in the SIA and DA.

To identify potential regulators essential for SIA and DA hematopoiesis, we compared their differentially expressed genes (DEGs), focusing on transcription factors (TFs). This revealed a number of candidate genes significantly enriched in either SIA or DA cells (Figure 6G). SIA-enriched TFs include *foxo4*, *hoxd3a* in SIA_ECs, *zfhx3b* and *nr4a3* in SIA_HECs, *gfi1ab* and *bcl11aa* in SIA_HSPCs (Figure 6G). Notably, *gfi1ab* and *bcl11aa* are well-documented in HSPC development and differentiation,^64–68^ while the roles of the other genes remain less explored. Similarly, DA-enriched TFs included *hoxb3a*, *mafba*, and *mafbb* in the DA_ECs, *sox18*, *ldb2a*, and *her4.4* in the DA_HECs, *znf296*, *nfil3-5*, and *nfil3-6* in the DA_HSPCs (Figure 6G). Among these, *hoxb3a*, *mafba*, *mafbb*, *sox18*, *nfil3-5*, and *nfil3-6* have been implicated in hematopoiesis,^69–73^ while the functions of the others are less studied. Together, these findings suggest that the distinct characteristics of SIA and DA hematopoiesis might be determined by specific sets of TFs.

## DISCUSSION

In this study, using time-lapse live imaging, short-term lineage tracing with photoconvertible fluorescent reporters, and permanent lineage tracing via the IR-LEGO cell labeling system, we documented that zebrafish HSCs originate from the subaortic SIA, rather than the DA. Furthermore, we identified bipotent VRHs in the SIA, capable of generating both hematopoietic and endothelial lineages. These findings challenge the conventional view of the DA as the primary source of HSCs and establish a new paradigm for studying HSC development and regulatory mechanisms.

An intriguing unanswered question is the source of the vascular progenitors that form the anterior SIA, which likely serve as precursors for HSC development. Previous studies suggest that the SIA endothelium is formed by two distinct vascular progenitors: ventral PCV ECs and cells derived from the extra-vascular secondary vascular field (SVF).^46–48^ Ventral PCV ECs sprout at approximately 28-56 hpf and migrate ventrally to contribute to the SIV and SIA endothelium,^46,47^ while SVF-derived cells, identified as *etv2^+^tal1^+^* vascular progenitor cells, emerge locally from the mesenchyme adjacent to the ventral PCV and integrate into the SIA and SIV endothelium.^48^ Given the minimal contribution of ventral PCV ECs to HSC production, as shown by lineage tracing in the PCV-marking *Tg(lyve1b:CreER^T2^)* line (Figures 2C and 2D), we postulate that SVF-derived *etv2^+^tal1^+^* cells are the likely precursors forming the anterior SIA endothelium, which then generate hematopoietic cells via EHT. Hence, developing an SVF-derived cell-specific CreER^T2^ line for fate mapping analysis will be essential to validate this hypothesis.

Another intriguing question is why the SIA generates HSCs while the DA primarily produces transient hematopoietic progenitor cells ― EHPs.^21^ One possible explanation is that the SIA and DA endothelium possess distinct intrinsic genetic/epigenetic features, dictating their differing hematopoietic potential. This hypothesis is supported by the scRNA-seq data, which reveals considerable transcriptomic differences between SIA and DA endothelial cells (Figures 6F and 6G). These distinct characteristics may be established early in development, as SIA and DA endothelium originate from different vascular precursor cells.^46–48^ Alternatively, extrinsic signals from the microenvironment may influence the hematopoietic potential of SIA- and DA-derived cells. For instance, as SIA is attached to the dorsal wall of the intestine, external cues from intestinal epithelial or smooth muscle cells could shape its hematopoietic traits. Similarly, the proximity of SVF cells and the SIA to pronephric duct during development^48^ suggest that epithelial cells of pronephros may create niches that influence HSC fate. The most plausible scenario likely involves a combination of intrinsic characteristics and external signals collectively regulating HSC fate in SIA-derived cells. Further investigation will be essential to elucidate these mechanisms.

Our study reveals a striking difference in hematopoietic cell production timelines between the DA and SIA. While hematopoietic cells are rapidly generated in the DA within two days through canonical EHT,^6^ the SIA employs both canonical and non-canonical EHT processes, sustaining hematopoietic cell production for at least two weeks. The coexistence of two EHT mechanisms suggests that the SIA may contain two distinct precursor types: unipotent HECs, which, akin to the HECs in the DA, produce hematopoietic cells through direct budding (canonical EHT), and bipotent VRHs, which undergo multiple rounds of division and budding, generating both hematopoietic and endothelial daughter cells (non-canonical EHT) (Figures 5D-5F; Movie S1). It remains unclear whether hematopoietic cells produced through canonical EHT and non-canonical EHT in the SIA differ in property and function. Given that the DA contains only unipotent HECs and primarily generates EHPs through canonical EHT,^6,40^ we speculate that SIA-derived hematopoietic cells produced via canonical EHT are likely to be EHPs, while those from non-canonical EHT are probably HSCs. The existence of bipotent VRHs provokes us to hypothesize that perhaps a similar cell type may exist in mammals. This hypothesis is supported by previous studies showing that the vascular marker VEGFR2 enriches human postnatal HSCs and that murine bone marrow HSCs can contribute to vascular endothelium upon transplantation.^74,75^ The zebrafish system offers a unique opportunity to explore the molecular characteristics, developmental regulations, and lifespan of the VRHs, providing insights into our understanding of hematopoietic and vascular system establishment and regeneration.

Finally, our research demonstrates that in zebrafish, HSCs primarily originate from the SIA, rather than the DA, challenging the traditional view of the DA as the primary source of HSCs in mammals. This discrepancy may arise from species differences, or alternatively, it reflects an evolutionary conserved mechanism, suggesting an alternative source for mammalian HSCs. We favor the latter possibility, in which two potential models can be suggested. The first model suggests that, similar to zebrafish, mammals may have an SIA-equivalent responsible for HSC production, while the DA primarily generates EHPs. Given that the SIA in zebrafish is a key embryonic artery dorsal to the intestine, we hypothesize that in mammals, embryonic intestinal arteries, particularly those dorsal to the intestine, including the vitelline artery and its derivatives like the celiac trunk and superior mesenteric artery,^76^ could serve as the major sources for HSC generation. This notion is supported by studies showing HSC activity in mouse vitelline artery during early embryogenesis.^12,77^ The second model suggests that, while mammals may lack a direct SIA equivalent, HSCs could arise from a unique group of subaortic mesenchyme-derived vascular precursors, similar to mesenchyme-derived *etv2^+^tal1^+^* endothelial precursors in zebrafish. In this scenario, these mesenchyme-derived vascular precursors may integrate with the DA endothelium post-DA formation, generating HSCs via EHT. Early studies demonstrating the presence of pre-HSCs in the subaortic mesenchyme of mouse embryos during or shortly after hematopoietic cluster formation in the DA support this idea.^29,30^ Recent work demonstrates that tamoxifen administration in *Tg(Pdgfra:CreER^T2^;ROSA26:LSL-eYFP)* mice at E9.5 labels a significant number of HSCs and HECs in the E11.5 DA.^78^ Given that *Pdgfra* is expressed in subaortic mesenchymal cells but not in the DA ECs, this finding further support the notion that mammalian HSCs may originate from subaortic mesenchyme-derived vascular precursors. Thus, a reevaluation of HSC origins in mammals is likely warranted.

## METHODS

### Zebrafish husbandry

All fish were maintained under standard conditions.^79^ The following transgenic lines were used in this study: *Tg(coro1a:loxP-DsRedx-loxP-eGFP)*,^22^ *Tg*(*kdrl:loxP-DsRedx-loxP-eGFP)^sz1^*^00^, *Tg(hsp70l:mCherry-T2A-CreER^T2^)*,^36^ *Tg(lyve1b:CreER^T2^)*,^80^ *Tg(stab1:DsRedx)^sz101^*, *Tg(kdrl:CreER^T2^)*,^81^ *Tg(fli1:eGFP)*,^82^ *Tg(fli1:loxP-DsRedx-loxP-eGFP)^sz102^*, *Tg(kdrl:eGFP)*,^83^ and *Tg(kdrl:Dendra2)*^21^.

### Generation of transgenic lines

The *kdrl:loxP-DsRedx-loxP-eGFP* plasmid was created by inserting the 6.4 kb *kdrl* promoter^83^ and the loxP-DsRedx-loxP-eGFP cassette^22^ into the pTol2 vector via Gibson assembly. Similarly, the *fli1:loxP-DsRedx-loxP-eGFP* plasmid was generated with a 2kb *fli1* enhancer-promoter fragment.^84^ A 2 kb promoter fragment upstream of the *stab1* ORF was amplified (FP: CTAAGGCTTTCTAGTTGGTTGT; RP: GTCTACTCCCTCAAAAGCCAG) from zebrafish genomic DNA and cloned into pTol2 vectors containing the DsRedx cassette, generating the *stab1:DsRedx* plasmid. To create transgenic lines, these plasmids (30 ng/μL) and in vitro transcribed transposase mRNA (50 ng/μL) were co-injected into one-cell-stage AB strain embryos, which were then raised to adulthood for germline transmission screening.

### Single-cell resolution & multi-cell resolution IR-LEGO lineage tracing

The experiments were conducted according to a previous protocol with minor modifications.^33^ Briefly, zebrafish embryos were anesthetized using 0.02% Tricaine (A5040; Sigma) and mounted in 1% low-melting agarose (A600015, Sangon Biotech). For single-cell resolution lineage tracing, targeted ECs/HECs were selectively heated by scanning a 1342 nm infrared light at 100 mW over a 6 mm x 8 mm area for 60 seconds to encompass a single cell. In multi-cell resolution lineage tracing, the infrared light was applied at 110 mW over an 8 mm x 8 mm area to heat targeted cells/tissues (e.g., the AGM and the SIA region). Each tissue received 3 to 4 sparse hits to label multiple cells. Following infrared light illumination (heat shock), embryos were treated with 10 μM (Z)-4-Hydroxytamoxifen (4-OHT) (H7904, Sigma) for 12-16 hours. After washing three times with egg water, the 4-OHT-treated embryos were raised and analyzed at the desired stages.

### Photoconversion assays

To trace the behaviors and distributions of SIA-derived cells, SIA HECs/ECs in 4 dpf *Tg(kdrl:Dendra2)* larvae were photoconverted using 405 nm laser with the photo-bleaching program on a Zeiss 980 or Leica SP8 confocal microscope.^85^ The photoconverted cells were immediately imaged to record the photoconversion pattern, and the distributions of their descendants were tracked for 1-4 days post-photoconversion. To characterize the dynamics of SIA HEC budding, a single SIA HEC/EC in the *Tg(kdrl:Dendra2)* zebrafish was photoconverted at various developmental stages (3-16 dpf). The photoconverted Dendra2-Red cells were subsequently tracked 24 hours post-photoconversion to determine the frequency of HEC budding.

### Lineage tracing with tissue-specific CreER^T2^ transgenic lines

To assess the contribution of PCV ECs or general vascular cells to adult HSCs, we crossed the venous/lymphatic vessel-specific *Tg(lyve1b:CreERT2)* fish or pan-vessel-specific *Tg(kdrl:CreER^T2^)* fish with the *Tg(kdrl:LRLG,coro1a:LRLG)* reporter line. The resulting triple transgenic embryos were treated with 4-OHT (10 μM) or vehicle (ethanol) from 12 hpf to 28 hpf to maximize labeling efficiency. At 3 dpf, 4-OHT-treated embryos were imaged to confirm PCV labeling efficiency. Each embryo with appropriate PCV or general vasculature labeling was raised to adulthood for flow cytometry analysis of adult blood cell labeling efficiency.

### Confocal imaging and time-lapse live imaging

All fluorescent images were captured with either a Zeiss 980 or Leica SP8 confocal microscope with 20x and 40x air objectives or a 40x water objective. All live embryos and larvae were mounted with 1% low-melting agarose in 0.5x E2 egg water. For time-lapse live imaging, we used either the aforementioned confocal microscopes or Olympus SpinSR spinning disc confocal microscope. Embryos and early larvae were mounted in 0.7% or 1% low-melting agarose and then covered with 0.02% Tricaine in 0.5x E2 egg water after the agarose solidified. Imaging dishes included 35 mm glass bottom confocal dishes (biosharp, BS-15-GJM) or µ-Slide 8 Well (ibidi, 80826), depending on the number of samples. Dishes were placed in a heating device to maintain a stable temperature of 28.5 ℃. Each imaging cycle was set for 15-20 minutes, and the samples were sequentially imaged over 12-48 hours to capture long-term cell behaviors. Images were processed with FIJI (ImageJ), except for Z-stack color-coding, which was processed via Zeiss ZEN 3.7 software.

### Single-cell suspension preparation and flow cytometry analysis

To assess the labeling efficiency of adult zebrafish KM blood cells, KM from euthanized adult zebrafish (3-12 months) were collected and mechanically dissociated in 400uL ice-cold 5% FBS/PBS buffer. The suspension was filtered through a 35 μm cell strainer (352235, Corning Falcon) to obtain a single-cell suspension for analysis. Flow cytometry analysis was performed using a BD FACSAria III flow cytometer, and data analysis, including gating image generation, was conducted using FlowJo (v10.8.1).

### Notch inhibitor treatment

The γ-secretase inhibitor DAPT powder (D5942, Sigma-Aldrich) was dissolved in DMSO to prepare a 100 mM stock solution. For experiments, 10 μL of the stock solution was mixed with 10 mL of 0.5x E2 embryo media to treat larvae aged 2.5-3.5 dpf, 3.5-4.5 dpf, and 2.5-4.5 dpf. In the control group, 10 µL of DMSO was mixed with the same volume of 0.5x E2 embryo media. After treatment, larvae were transferred to regular 0.5x E2 embryo media or system water for further development until subsequent assays.

### Manual cell picking and Smart-seq2 scRNA-seq sample processing

To prepare single-cell suspension for manual cell picking, 5 dpf *Tg(kdrl:Dendra2)* larvae with photoconverted anterior SIA were anesthetized using 0.02% Tricaine (E10521, Sigma-Aldrich). Trunk containing photoconverted SIA was dissected from ∼80-100 larvae, washed with ice-cold 5% FBS/PBS buffer, and mechanically disassociated using 26 G needles and syringes. The dissociated tissues were enzymatically digested with EDTA-free TrypLE^TM^ Express enzyme (12605028, Gibco) for 10 min, followed by 1:1 mixed TrypLE^TM^ Express and StemPro^TM^ Accutase^TM^ (A1110501, Gibco) for an additional 10 min. The digestion reaction was stopped by adding FBS to a final concentration of 10%. Digested cells were harvested by standard centrifugation, washed, and resuspended in ice-cold 1% BSA/DPBS (14190144, Gibco) buffer. Finally, the cell suspension was filtered through a 35 μm cell strainer to obtain a single-cell suspension.

To pick photoconverted cells, 20 μL of the single-cell suspension was carefully loaded into an uncoated sterile culture dish. Individual photoconverted cells were collected using an IM-9C pneumatic injector (Narishige) under a fluorescent microscope and immediately transferred into sterile 200 μL PCR tubes preloaded with 4.4 μL lysis buffer. Immediately following the transfer, the cells were vortexed, spun down, and stored at -80°C.

Whole-transcriptome amplification and library preparation were performed following the Smart-seq2 protocol.^86,87^ Library quality was evaluated using either the Agilent 5300 Fragment Analyzer or the BiOptic Qsep100 Fragment Analyzer. Prepared libraries were sent to Novogene for Illumina 150-bp paired-end sequencing, with an average sequencing depth of 2 Gb per sample.

### scRNA-seq data integration and analysis

For scRNA-seq data analysis, reads were aligned to the GRCz11.113 zebrafish reference genome using the STAR package (2.5.2b),^88^ and gene counts were quantified with featureCounts (Rsubread_2.20.0). The resulting count matrices were normalized to transcripts per million (TPM) for downstream analysis. Datasets from 5 dpf SIA and 28 hpf-DA photoconverted cells were processed and analyzed using the Seurat package (v.5.1.0).

Prior to analysis, quality control was performed using the following criteria: nCount_RNA > 200, nFeature_RNA < 2000, and percent.mt < 5. After filtering, 181 cells were retained for analysis. Principal component analysis (PCA) was performed, followed by data scaling. To minimize batch effect, the Harmony algorithm was applied as part of the standard integration workflow (https://satijalab.org/seurat/articles/seurat5_integration). Clustering analysis was conducted, and visualization was accomplished through dimensionality reduction using UMAP based on shared nearest-neighbor clustering. Dot plots, feature plots, and violin plots were generated to visualize gene expression patterns using SCT-normalized data. GO enrichment analysis was performed with the clusterProfiler package (v.4.14.4), using marker genes filtered by a minimum percentage of cells expressing the gene (min.pct = 0.25) and a log fold change threshold (logfc.threshold = 0.25). All analyses were conducted within the R environment (v.4.4.2).

### Quantifications and statistics

Statistical parameters, including the exact value of n, are reported in the figures and figure legends. Data are presented as means ± SD whenever applicable. Statistical significance is denoted as follows: ns, P>0.05; * P ≤ 0.05; ** P ≤ 0.01; *** P ≤ 0.001, and **** P ≤ 0.0001. Depending on variance homogeneity, unpaired Student’s t-test or Welch’s t-test were used to calculate P values. All statistical analyses were performed using GraphPad Prism version 10.

## Supporting information

Movie S1_No EHT in PCV

Movie S2_EHT process of an SIA cell

Movie S3_No obvious EHT in SIA in runx1 mutant fish

## ACKNOWLEDGEMENTS

This work was supported by grants from the National Key Research and Development Program of China (2023YFA1800100), the Shenzhen Medical Research Fund (B2302034), and the Research Grants Council of Hong Kong (T13-602/21-N).

## AUTHOR CONTRIBUTIONS

Z.W. and S.F. conceived the project, designed the experiments, and wrote the manuscript. S.F. and R.Q. performed all the experiments. S.H., Y.H., and T.H. provided the single-cell resolution IR-LEGO system and assisted with lineage tracing experiments. Y.W., T.T., K.C., S.Z., Y.H., and H.Z provided additional experimental support.

