## Supplementary files for "Bona fide hematopoietic stem cells in zebrafish originate from the supra-intestinal artery"

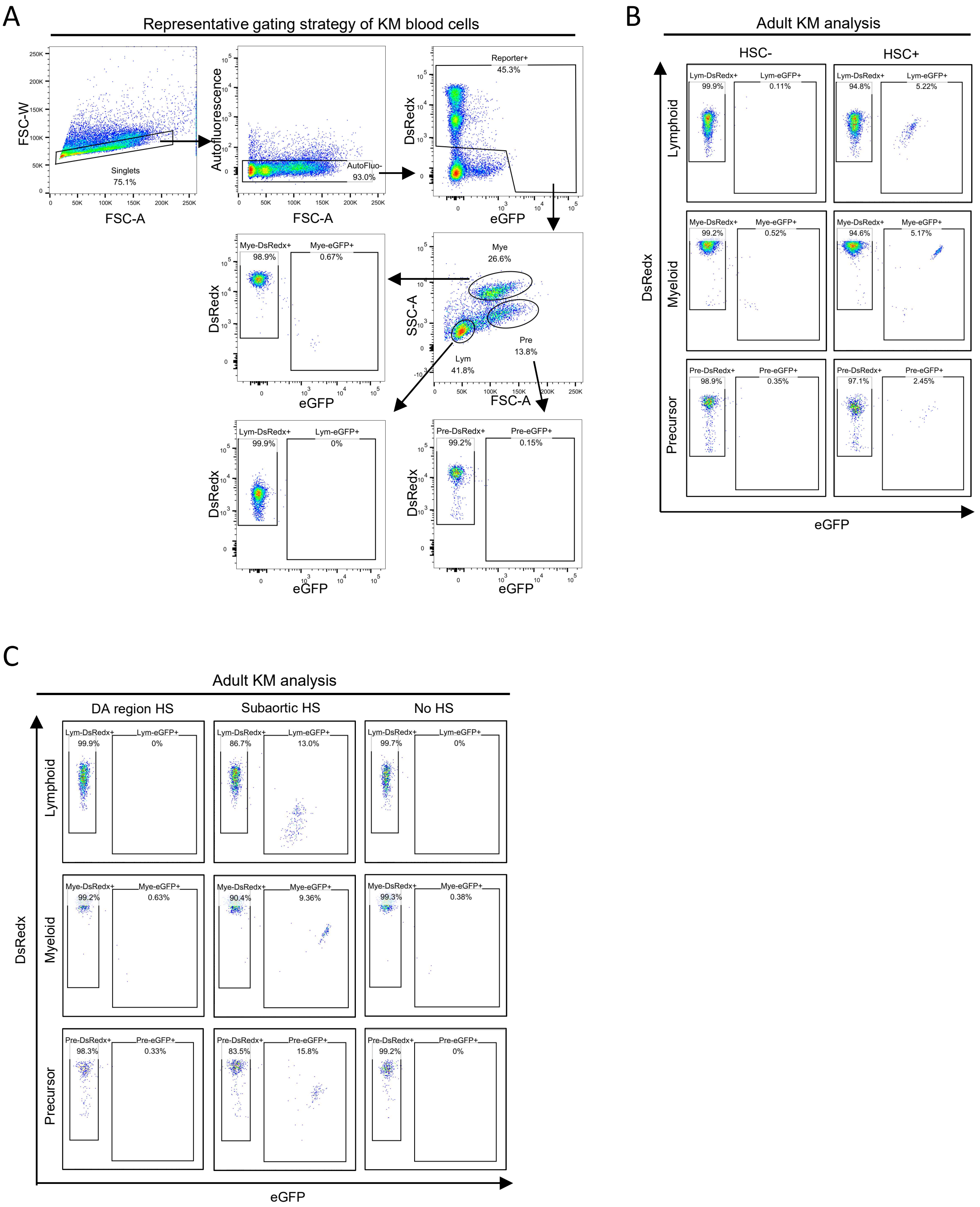


**Figure S1. Flow cytometry analysis of adult kidney marrow blood cells, related to Figure 1**

(A) Related to Figure 1C and all KM blood cell analysis experiments. Schematic illustration of flow cytometry gating strategy for adult KM blood cells. KM: kidney marrow.

(B) Related to Figure 1C. Representative flow cytometry analysis of hematopoietic cells (eGFP$^+$) in adult KM from HSC-labeled (HSC+) and HSC-unlabeled (HSC-) control fish. Fish with ≥1% eGFP$^+$ hematopoietic cells are classified as HSC+, while those with <1% eGFP$^+$ hematopoietic cells are defined as HSC-.

(C) Related to Figure 1F and 1G. Representative flow cytometry analysis of adult KM blood cells from DA region-labeled fish, Subaortic AGM region-labeled fish, and No HS (control) fish.

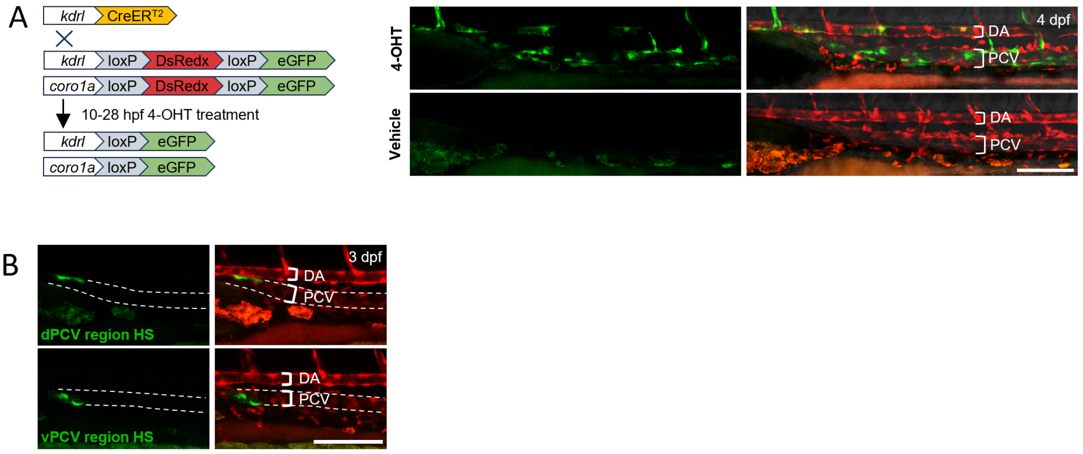


**Figure S2. HSCs originate from early *kdrl*$^{+}$ cells in the AGM subvenous region, related to Figure 2**

(A) Related to Figure 2C and 2D. Schematic illustration of *kdrl*:CreER$^{T2}$-mediated lineage tracing. The left panel shows the principle of the experiment, and the right panel indicates the AGM vasculature labeling pattern.

(B) Representative images of dPCV and vPCV region-labeled fish. Fish with relatively region-specific labeling were selected based on the eGFP$^{+}$ vasculature pattern observed at 3 dpf using confocal microscopy.

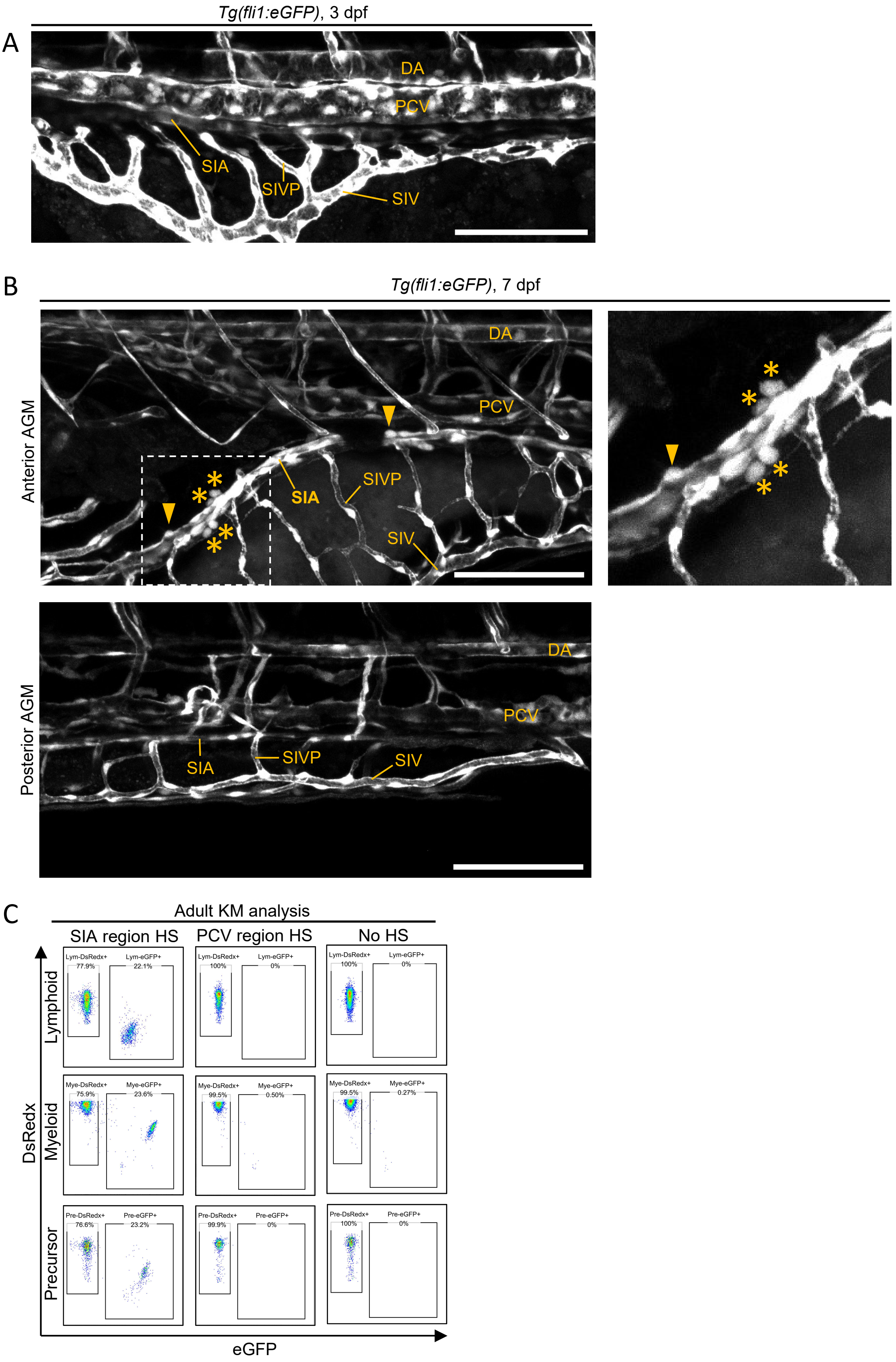


**Figure S3. Characterization of hemogenic potential of the subvenous vasculatures, related to Figure 3**

(A) A representative image of subvenous vasculatures in *Tg(fli1:eGFP)* fish during 2-

4 dpf. EHT-like events or eGFP$^+$ nascent HSPC-like cells were barely detected. SIA: supra-intestinal artery. SIV: sub-intestinal vein. SIVP: sub-intestinal venous plexus.

(B) High resolution images of the subvenous vasculatures reveal EHT-like events (arrowhead) in the SIA and eGFP$^+$ nascent HSPC-like cells derived from the SIA (asterisks) (upper panel) in the anterior AGM region of 7 dpf *Tg(kdrl:eGFP)* fish. The boxed region is magnified in the upper right panel. In contrast, no EHT-like events or eGFP$^+$ nascent HSPC-like cells were detected in the posterior AGM region (lower panel)

(C) Related to Figure 3F and 3G. Representative flow cytometry analysis of adult KM blood cells of SIA region-labeled fish, PCV region-labeled fish, and No HS (control) fish.

### Supplementary movies information

**Movie S1, related to Figure 2.**

Time-lapse live imaging of the anterior PCV during 29 hpf to 72 hpf in *Tg(stab1:DsRedx)* fish. No EHT-like process is detected. Some autofluorescence-positive pigment cells autonomously emerge in the subvenous region.

**Movie S2, related to Figure 3.**

Time-lapse live imaging of the subvenous vasculatures in 4-6 dpf *Tg(kdrl:eGFP)* fish reveals EHT-like processes in the anterior SIA. Each frame of the video represents a single slice from a z-stack, providing optimal details of the EHT process. The frequent autonomous contractions of the intestine cause movement of the SIA.

**Movie S3, related to Figure 4.**

Time-lapse live imaging of the anterior SIA in 5-5.5 dpf *runx1*$^{-/-}$ *Tg(kdrl:eGFP)* fish shows a reduction in EHT-like processes. Each frame is a maximum projection of a z-stack. Frequent autonomous contractions of the intestine cause SIA distortion, affecting the quality of some frames.
